# Multi-SNP Mediation Intersection-Union Test

**DOI:** 10.1101/455352

**Authors:** Wujuan Zhong, Cassandra N. Spracklen, Karen L. Mohlke, Xiaojing Zheng, Jason Fine, Yun Li

## Abstract

Tens of thousands of reproducibly identified GWAS (Genome-Wide Association Studies) variants, with the vast majority falling in non-coding regions resulting in no eventual protein products, call urgently for mechanistic interpretations. Although numerous methods exist, there are few, if any methods, for simultaneously testing the mediation effects of multiple correlated SNPs via some mediator (for example, the expression of a gene in the neighborhood) on phenotypic outcome. We propose SMUT, multi-<u>S</u>NP <u>M</u>ediation intersection-<u>U</u>nion <u>T</u>est to fill in this methodological gap. Our extensive simulations demonstrate the validity of SMUT as well as substantial, up to 92%, power gains over alternative methods. In addition, SMUT confirmed known mediators in a real dataset of Finns for plasma adiponectin level, which were missed by many alternative methods. We believe SMUT will become a useful tool to generate mechanistic hypotheses underlying GWAS variants, facilitating functional follow-up. The R package SMUT is publicly available from CRAN at https://CRAN.R-project.org/package=SMUT.

## INTRODUCTION

Genome-wide association studies (GWASs) have been successful for detecting genetic variants associated with complex diseases and traits, which usually result from interplay of multiple factors including genetic, epigenetic, transcriptomic, proteomic and/or environmental factors. The effects of genetic variants either individually or even in aggregation on complex traits are typically small to moderate at best (1). More importantly, the vast majority of identified GWAS variants (in the order of >10^4^) do not map to protein-encoding regions, so the underlying mechanism remain largely elusive. Expression quantitative trait loci (eQTL) analysis which postulate mechanistic hypotheses between genetic variants and the expression levels of genes, particularly genes in the neighborhood (i.e., *cis* or more precisely local eQTLs) (2–7), has become an important tool for functional interpretation of GWAS. Transcriptome-wide association studies (TWAS), which identify significant expression-trait associations through correlating the imputed gene expression to the trait, enables the GWAS and eQTL datasets from two independent studies (8–10, BioRxiv: https://doi.org/10.1101/286013). Such TWAS analyses can also be performed using summary statistic from GWAS and eQTL datasets when individual level data are not available (11). Mancuso et al. proposed a method of utilizing the *cis* genetic variation near a gene to estimate the local genetic correlation between gene expression and a complex trait in TWAS and estimate the causal relationship between pairs of complex traits that are genetically correlated (11). Integration of genotype, gene expression and phenotype information from GWAS and eQTL datasets will fundamentally advance our knowledge of molecular mechanisms of complex disorders and quantitative traits. Several excellent review papers exist for causal relationship inference in the context of genetic mapping for complex traits (12, 13).

The integrative genomic studies enable mechanistic interpretations, for example, via either the methods of instrumental variable(s)(IV[s]) and/or mediation analysis. Mendelian randomization (MR) framework (14–16) has been adopted by a number of methods. MR treats genetic variant(s) (in most cases, SNPs) as the IV(s) to assess the causal effects of genetic variants through some mediator(s) of interest (e.g. expression levels of some gene[s]) on the trait of interest (10, 15, 17). Classic MR methods, such as SMR (10), make several key assumptions including complete mediation, where SNPs must be marginally independent of the confounding between mediator and final outcome, and a priori knowledge that the causal flow is from SNP to mediator but not the reverse (14, 18, 19). When the assumptions are violated, MR performs essentially invalid IV analysis, leading to biased inference. Some of the more recently developed MR methods allow relaxation of certain key assumption(s) aforementioned. Such relaxation(s), however, are at costs. For example, MR-Egger (20, 21) relaxed the complete mediation assumption and allows multiple IVs by first analyzing each IV individually and then meta-analyzing individual IV results. However, MR-Egger assumes that multiple IVs (i.e., SNPs) in the analysis are uncorrelated, limiting its application to a typical GWAS or eQTL locus where multiple partially dependent SNPs are identified.

Another drawback of MR methods is that they cannot distinguish between mediation and pleiotropy, the phenomenon of one SNP having effect on more than one outcome (where the outcome can be either a molecular measure such as gene expression or a phenotypic outcome) (Figure 1). Since pleiotropy is commonly observed for many complex traits (22), MR methods are therefore not preferred, when the goal is to infer mediation or to generate mechanistic hypotheses. The more recent CaMMEL method (BioRxiv: https://doi.org/10.1101/219428) further extends MR-Egger to allow multiple mediators and to model multiple IVs simultaneously (in contrast to MR-Egger which models each IV individually and then meta-analyzes individual IV results). In addition, CaMMEL accommodates linkage disequilibrium (LD) among SNPs (23). Thus, CaMMEL can handle multiple correlated genetic variants and claims to be more powerful than Mancuso et al. method (11) and MR-Egger (24) without inflated false discovery rates (FDR). However, CaMMEL, designed for multiple mediators modeled simultaneously, is sub-optimal for single mediator analysis. In addition, CaMMEL assumes the presence of at least one eQTL since it tests the effect of mediators on phenotype (as in contrast to testing the presence of indirect genotype effect via mediator(s) on phenotype).

**Figure 1.**
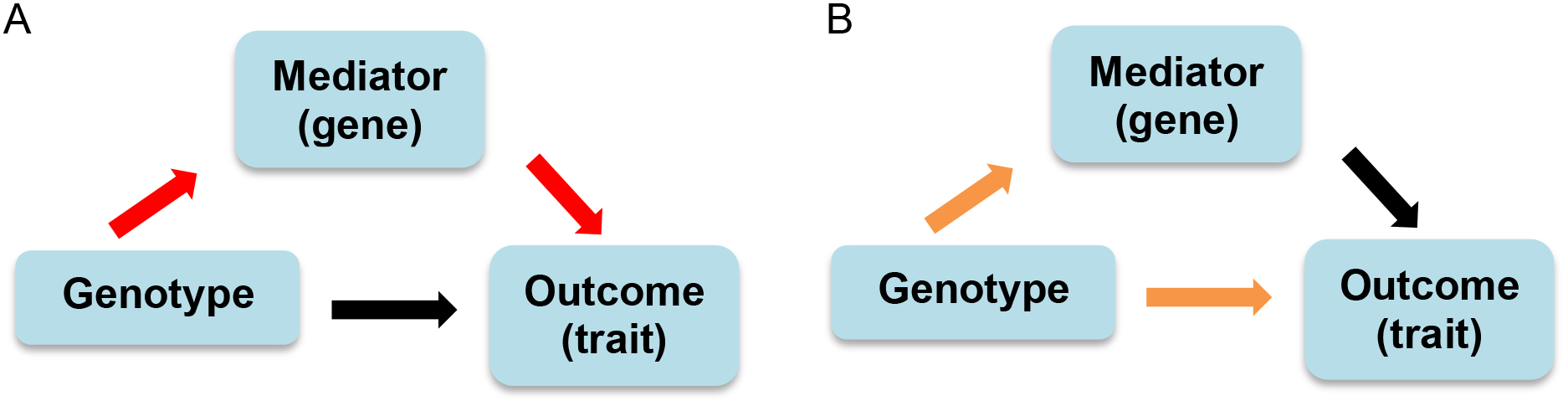
Directed acyclic graph for mediation and pleiotropy. (**A**) Red arrows indicate the mediation effect of the genotype on the outcome through the mediator. (**B**) Orange arrows indicate the pleiotropy.

Besides TWAS and MR, other mechanism elucidating methods proposed in the recent literature include CIT (Causal Inference Test) (25) and Huang et al. (26–28). CIT employs regression based framework and tests for complete mediation and is thus most suitable under the arguably unrealistic scenario of complete mediation. The methods of Huang et al. employ a kernel regression framework and uses variance component score statistic (29) to test for mediation. However, this method also assumes that genetic variants under testing are, or at least contain, *a priori* known eQTL(s). In addition, it only tests the effect of mediators on phenotype, similar to CaMMEL, again in contrast to testing indirect genotype effect through mediators on phenotype.

Popular approaches of classic mediation analysis include causal steps, difference method and product method (30, 31). The causal steps approach performs multiple tests involved in a causal chain. The difference method is based on the difference in the coefficient estimate of the treatment (here, genetic variant) before and after including the mediator in the regression model. The product method, such as the Sobel test (32, 33), explicitly tests the product of the treatment coefficient in the mediator model and the mediator coefficient in the outcome model. However, it is unclear that such methods can be adapted to integrative genomic settings with high dimensional SNPs.

In short, to the best of our knowledge, few, if any, existing methods can simultaneously accommodate incomplete mediation as well as multiple correlated SNPs, when complete individual level data (including genotype, mediator, and phenotype information) are available. To fill the gap, we propose here SMUT, multi-SNP Mediation intersection-Union Test, to explicitly accommodate both direct and indirect (via mediator) effect of multiple (in the order of hundreds to thousands) correlated SNPs on phenotype of interest. SMUT is a flexible, regression based approach that evaluates the joint mediation effects of multiple genetic variants on some trait of interest through a single mediator. SMUT extends the classic framework of Baron and Kenny (34) to allow multiple treatment variables (in our context, multiple genetic variants). Leveraging the intersection-union test (IUT) (35), SMUT decomposes mediation effects using two separate regression models. One is the mediator model where we regress the mediator on multiple genetic variants. For this mediator model, SMUT adopts the SKAT (36, 37) framework to handle a potentially large number of genetic variants in a statistically and computationally efficient manner. The other is the outcome model where we regress the outcome on both the mediator and multiple genetic variants. Classic regression models fail for the outcome model due to the high dimensionality of the SNPs. To solve this issue in SMUT’s outcome model, we adopt a mixed effects model and the Rao’s score test (38, 39) for mediation testing. Our extensive simulations and real data analysis demonstrate the advantages of SMUT over alternative methods. For example, with controlled type-I error, we show up to 92% power gain in simulations. More importantly, in real data analysis, SMUT confirms mediations at several well-established positive control loci while most of the alternative methods failed to reveal any of the relationships.

## MATERIAL AND METHODS

### multi-SNP Mediation intersection-Union Test (SMUT)

SMUT is a powerful test for the joint mediation effects of multiple genetic variants on a trait through a single mediator. The multiple genetic variants can be in a region, sub-locus defined by genes, or moving windows across the genome.

### Notation and Data Set-up

Without loss of generality, we assume that we have three types of data. Specifically, genotypes, gene expression measurements (can be other types of mediators such as metabolite levels or protein abundances) and phenotypic trait are available. Let *G* = (*G*_1_, *G*_2_, …, *G_q_*) be the *n* by *q* genotype matrix, where *n* is sample size, *q* is the total number of marker, and *G_j_* = (*G*_1*j*_, *G*_2*j*_,…, *G*_*nj*_)*^T^* is the vector of genotypes for the *n* samples at marker *j*, *j* = 1, 2, …, *q*. We consider an additive model with *G_ij_* taking values 0,1,2, measuring the number of copies of the minor allele. Suppose in total there are *l* genes *M*, *M*^(2)^, *M*^(3)^, …, *M*^(l)^ with the first notation *M* having no superscript. Here, *M* = (*M*_1_, *M*_2_, …, *M*_*n*_)*^T^*, is the vector of expression values of a given gene (the mediator) for *n* samples. Similarly, *M*^(2)^, …, *M*^(*l*)^ are the vectors of expression values of the other (*l* – 1) genes (i.e., mediators). Let *Y* = (*Y*_1_, *Y*_2_, …, *Y*_*n*_)^*T*^ be the vector of phenotypic trait.

### SMUT Model and Test for Joint Mediation Effects

SMUT models the effects of genetic variants on the trait mediated by the expression level of a single gene. We start with considering a more general model with multiple genes expression levels via the following regression models:

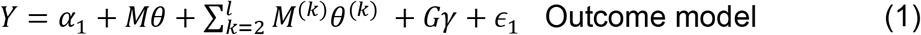

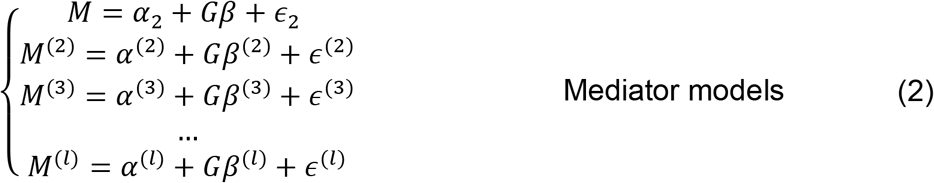

where *γ* = (*γ*_1_, *γ*_2_, …, *γ_q_*)*^T^* are the direct effects of the *q* genetic variants; *βθ* measures the indirect effects mediated by *M* for the multiple genetic variants. Similarly *β*^(*k*)^*θ*^(*k*)^ measures the indirect effects mediated by *M*^(*k*)^, *k* = 2,3,…, *l*.

Substituting the *M*, *M*^(2)^, *M*^(3)^, …, *M*^(*l*)^ with the values in (2), we have

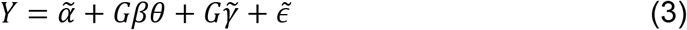

Where 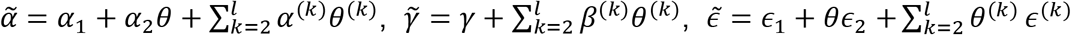

Equation (3) shows that indirect effects mediated by *M*^(2)^, *M*^(3)^, …, *M*^(*l*)^ would be absorbed by the direct effects 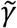 if we only model gene *M*. Therefore, without loss of generality, we only consider the mediation analysis for a given single gene expression level and consider the regression models below

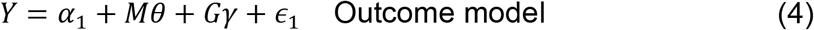

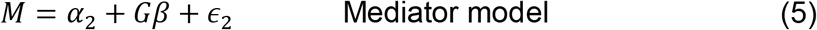

where 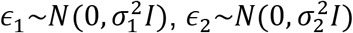, and we assume that *ϵ*_1_ and *ϵ*_2_ are independent; otherwise their correlation would make themselves mediator-outcome confounders which violates the key assumption for mediation analysis (30, 31).

Here *γ* measures effects from two sources: direct effects of the *q* genetic variants on outcome; and indirect effects of genetic variants via mediators other than *M*. For presentation brevity and clarity, we hereafter use direct effects to refer to the aggregated effects from the above two sources. We are interested in testing the mediation effect, of the *q* genetic variants via mediator *M*. Specifically, we test the null hypothesis H_0_: *βθ* = 0. If we have only one genetic variant, then *βθ* would be a scalar and the classic methods for testing mediation effects, such as the Sobel test (32, 33), under the framework of Baron and Kenny can be applied. Since we focus on the joint (from multiple genetic variants) mediation effects, *βθ* is thus a vector in our setup. The null hypothesis again is H_0_: *βθ* = 0, versus the alternative hypothesis H_1_: *βθ* ≠ 0. The null hypothesis is divided into two sub-hypotheses, 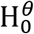: *θ* = 0 versus 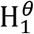: *θ* ≠ 0 and 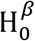: *β* = 0 versus 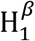: *β* ≠ 0. Thus, we have

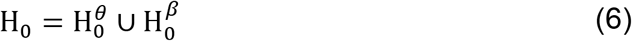

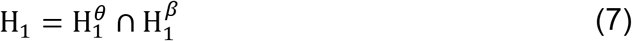

This can be conveniently solved by the intersection-union test (IUT). Suppose the *p* value for testing 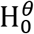 versus 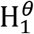 is *p*_1_; and the *p* value for testing 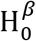 versus 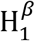 is *p*_2_. Then the *p* value for testing the overall H_0_ versus H_2_ applying IUT is the maximum of *p*_1_ and *p*_2_. In the following sections, we use the SMUT strategy to test *θ* and *β* separately to obtain and *p*_2_.

### Testing β in the Mediator Model

Many of the testing methods for association between multiple genetic variants and the trait can be applied here. We adopt the SKAT framework, a de facto locally powerful test(36), which accommodates large numbers of genetic variants efficiently.

### Testing θ in the Outcome Model

The outcome model is also high dimensional with multiple genetic effects and the mediator. Classic regression models tend to fail for such models. As a solution, we employ the following mixed effects model to reduce the dimension of parameters.

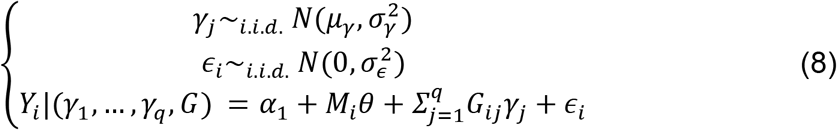

We first write out the log-likelihood function for model (8) and then derive the Rao’s score statistic (38, 39) for testing *θ*. Next, we apply Expectation–maximization (EM) algorithm to obtain maximum likelihood estimate (MLE) under the null hypothesis (40, 41). Finally, the score statistic is evaluated at MLE.

The log-likelihood for outcome *Y* is

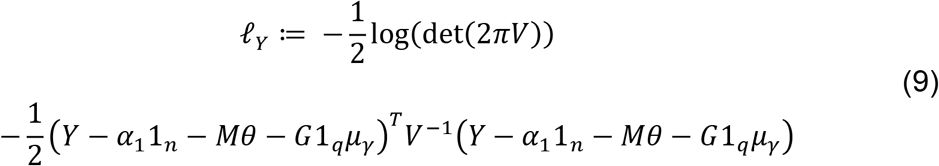

where 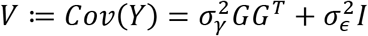 and 1_*k*_ := (1,1,…,1)^*T*^ is a vector of *k* copies of 1.

The Rao’s score statistic for testing *θ* is

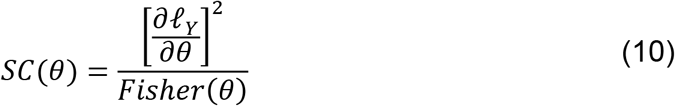

where 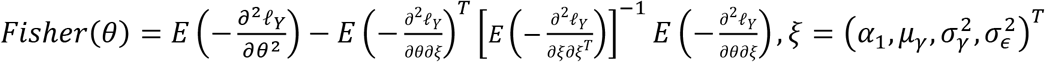

Derivations can be found in (41). The first and second derivatives of *ℓ_Y_* for our model are detailed in the Supplementary Data.

Under the null hypothesis *θ* = 0, this score statistic *SC*(*θ*) asymptotically follows a Chi-squared distribution with one degree of freedom when MLE under the null is plugged in. This assumes at least some of the direct effects *γ_j_*(*j* = 1,2,…,*q*) are nonzero. When there are no direct effects, the variance component 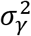 is on the boundary. The asymptotic Chi-square distribution works well in simulations (Supplementary Figure S1 and S2).

We leverage the EM algorithm to obtain MLE under the null. When applying EM algorithm to mixed effects model, random effects *γ* are treated as missing data. The complete data comprise the observed outcome data and random effects. The log-likelihood for complete data (*Y, γ*) is

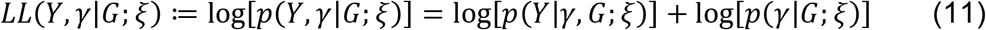

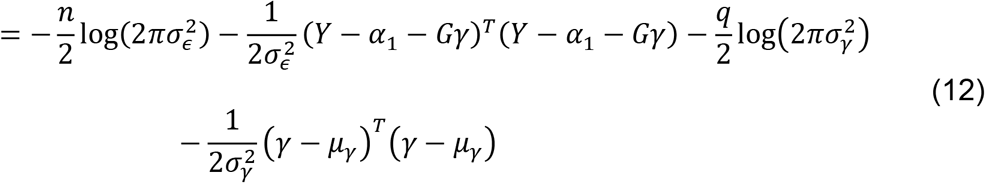

where 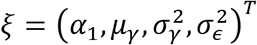

Derivations for E-step and M-step can be found in (41).

E-step of EM algorithm is

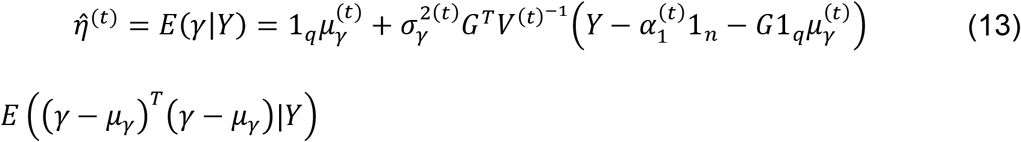

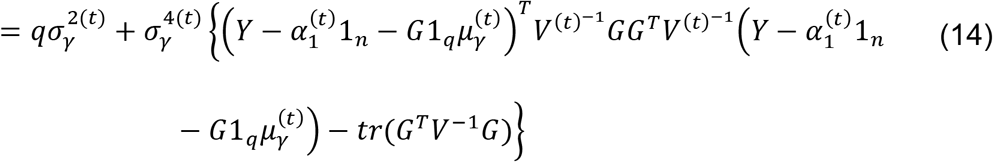

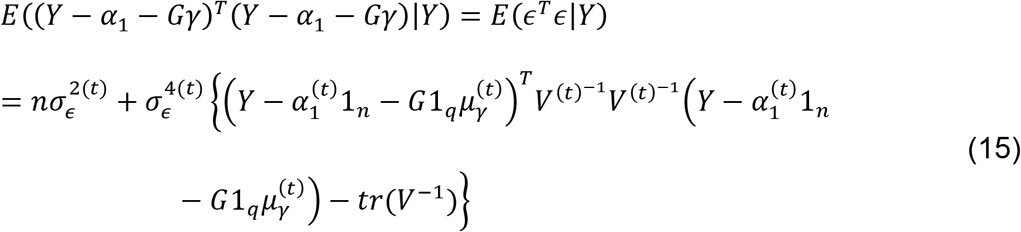

M-step of EM algorithm is

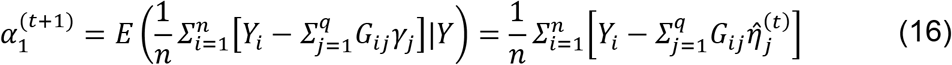

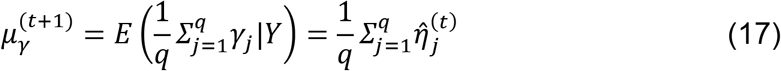

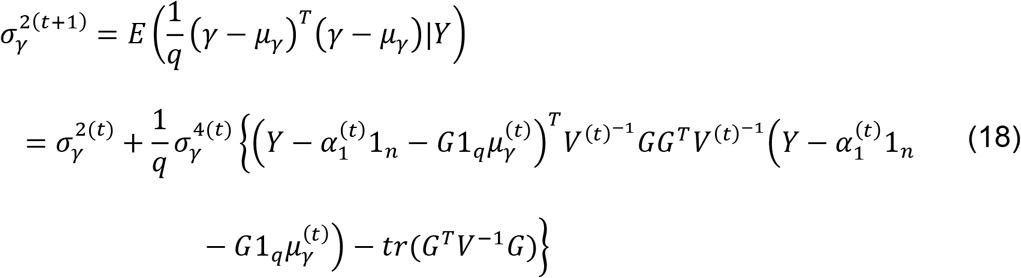

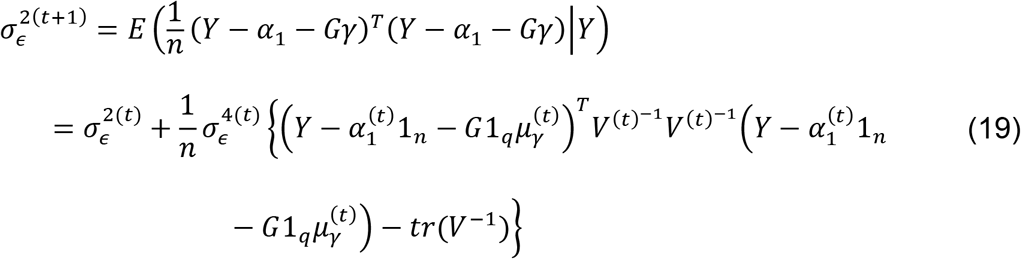

Convergence criterion for EM algorithm is

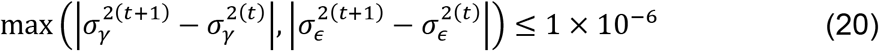

If convergence is not reached, iteration stops when the number of iterations exceeds a pre-specified large number.

As for the starting values of EM algorithm, the intercept *α*_1_ is randomly generated from uniform distribution *Unif*(−1,1). And *μ_γ_* is also randomly generated from uniform distribution *Unif*(−1,1). The variance components 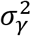 and 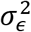 are independently generated from uniform distribution *Unif*(0,1).

### Simulations

To evaluate the performance of SMUT in comparison with alternative methods, we carried out extensive simulations to investigate power and type-I error. We first simulated 20,000 European-like chromosomes in a 1Mb region, using the COSI coalescent model (42) to generate realistic data in terms of allele frequency, linkage disequilibrium and population differentiation. The final dataset had 23,889 SNPs in a 1 Mb region. We constructed 10,000 pseudo-individuals by pairing up the 20,000 simulated chromosomes. To evaluate power and type-I error, we generated 200 datasets with 1,000 samples each by sampling without replacement from the entire pool of 10,000 samples above. Simulations were restricted to the 2,891 SNPs with minor allele frequency (MAF) ≥ 1%.

The outcome (trait) and the mediator were generated via the following outcome model (21) and mediator model (22), respectively.

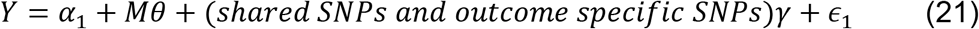

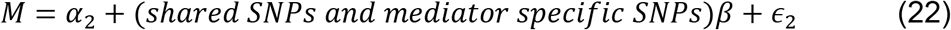

Where *β* ~ *c_β_N*(2,2),*γ* ~ *c_γ_N*(2,2),*ϵ*_1_ ~ *N*(0,1),*ϵ*_2_ ~ *N*(0,1).

We set *c_γ_* = 0.2 to evaluate the performance of SMUT and alternative methods under the scenario of pleiotropy. Specifically, the shared SNPs (sSNPs) between the two models are those that influence both the mediator and the outcome trait. The outcome (or mediator) specific SNPs only contribute to the trait (or mediator). The causal SNPs are the union of the sSNPs, mediator specific SNPs (mSNPs) and outcome specific SNPs (oSNPs). We considered two scenarios in terms of causal SNP density: sparse and dense (Table 1), with 10 and 1,000 causal SNPs respectively. The set of (10 or 1000) causal SNPs, common across the 200 datasets, were randomly selected from the 2,891 SNPs with MAF ≥ 1%. *β* and *γ*, again fixed across the 200 datasets, were independently drawn from a normal distribution with mean and variance both being 2.

**Table 1.** Causal SNP composition in two simulated scenarios. The sparse(dense) scenario is to simulate data sets based on a small(large) number of causal SNPs. Causal SNPs are the union of shared SNPs, mediator specific SNPs and outcome specific SNPs. Shared SNPs have effects on both mediator and outcome. Mediator(outcome) specific SNPs have effects only on mediator(outcome). All these SNPs are randomly selected from the 2,891 SNPs with MAF ≥ 1%.

|  | # of causal SNPs | # of shared SNPs | # of mediator specific SNPs | # of outcome specific SNPs |
| --- | --- | --- | --- | --- |
| **Sparse** | 10 | 4 | 3 | 3 |
| **Dense** | 1000 | 334 | 333 | 333 |

Error terms *ϵ*_1_ and *ϵ*_2_ were independently generated from standard normal distribution and were separately simulated for each of the 200 datasets.

In the simulations, we tested the joint mediation effects of these 2,891 SNPs on the trait using SMUT and other methods including adaptive Huang et al.’s method, adaptive LASSO (43), adaptive CaMMEL, Sobel test, and SMR. The adaptive Huang et al.’s method adopts the original kernel framework where effect(s) of interest are treated as random (26–28). In our context, when applying Huang et al., we treat the mediator coefficient in the outcome model as a random effect and apply IUT using variance component score test in the outcome model and SKAT in the mediator model. The adaptive LASSO employs LASSO for variable selection in the outcome model and applies IUT using regular regression with the selected variables in the outcome model and all the variables (i.e., genetic variants) via SKAT framework in the mediator model. The adaptive CaMMEL applies IUT using CaMMEL to test *θ* in the outcome model and SKAT to test *β* in the mediator model. Since Sobel test and SMR can only model one single SNP at a time, we tested each SNP separately and applied Bonferroni adjustment.

As detailed above, we simulated causal SNPs only from the pool of common (MAF > 1%) SNPs. By default, we tested all common SNPs in the region to mimic the realistic scenario where we have relatively little information regarding which SNPs are causal, at an established GWAS locus. To test the robustness and generalizability of the methods, we considered two alternative testing strategies each with a reduced set of genetic variants modelled. For the first testing strategy, we assume prior knowledge of eQTL SNPs (union of shared and mediator specific causal SNPs) and test only these eQTL SNPs. On the positive side, such an approach results in a reduced model with causal SNPs considered only. On the negative side, a subset of causal markers (specifically, the outcome specific causal SNPs) are not modelled. The second strategy tests SNPs with MAF ≥ 5%, thus missing true causal SNPs with MAF between 1% to 5%.

## RESULTS

### Type-I Error in Simulations

We evaluated SMUT along with alternative methods in simulations. SMUT manifested controlled type-I error rates, at *α* = 0.05 level, regardless of causal SNP density, as shown in Figures 2 and 3 for sparse and dense scenarios, respectively. Note that the first panel (*c_β_* = 0) and the leftmost point (*θ* = 0) in other panels (*c_β_* ≠ 0) all correspond to the null of no mediation through the mediator. Adaptive Huang et al.’s method also showed protected Type-I error. In contrast, Sobel test and SMR showed substantial inflation in Type-I error, particularly when *c_β_* is large. For example, when *c_β_* = 0.2, *θ* = 0 and sparse causal SNPs, Type-I error rates for Sobel test and SMR are 90% and 100% respectively. Such marked inflation in Type-I error is likely due to the more severe violation of the assumption of no pleiotropy, made by these two methods, as *c_β_* increases. Adaptive CaMMEL also showed Type-I error inflation. For example, among sparse causal SNPs when *c_β_* = 0.2, *θ* = 0, the Type-I error rate is 100%. We suspect such inflation is due to the fact that CaMMEL was developed for joint testing of multiple mediators via a Bayesian framework to borrow information across mediators. Thus, when testing one single mediator, lack of information in the Bayesian inference can lead to Type-I error inflation. Adaptive LASSO had severe Type-I error inflation when the causal SNPs were dense (Figure 3). For instance, when *c_β_* = 0.05 and *θ* = 0 type-I error rate is 75%. This is likely due to the violation of LASSO’s sparsity assumption (44).

**Figure 2.**
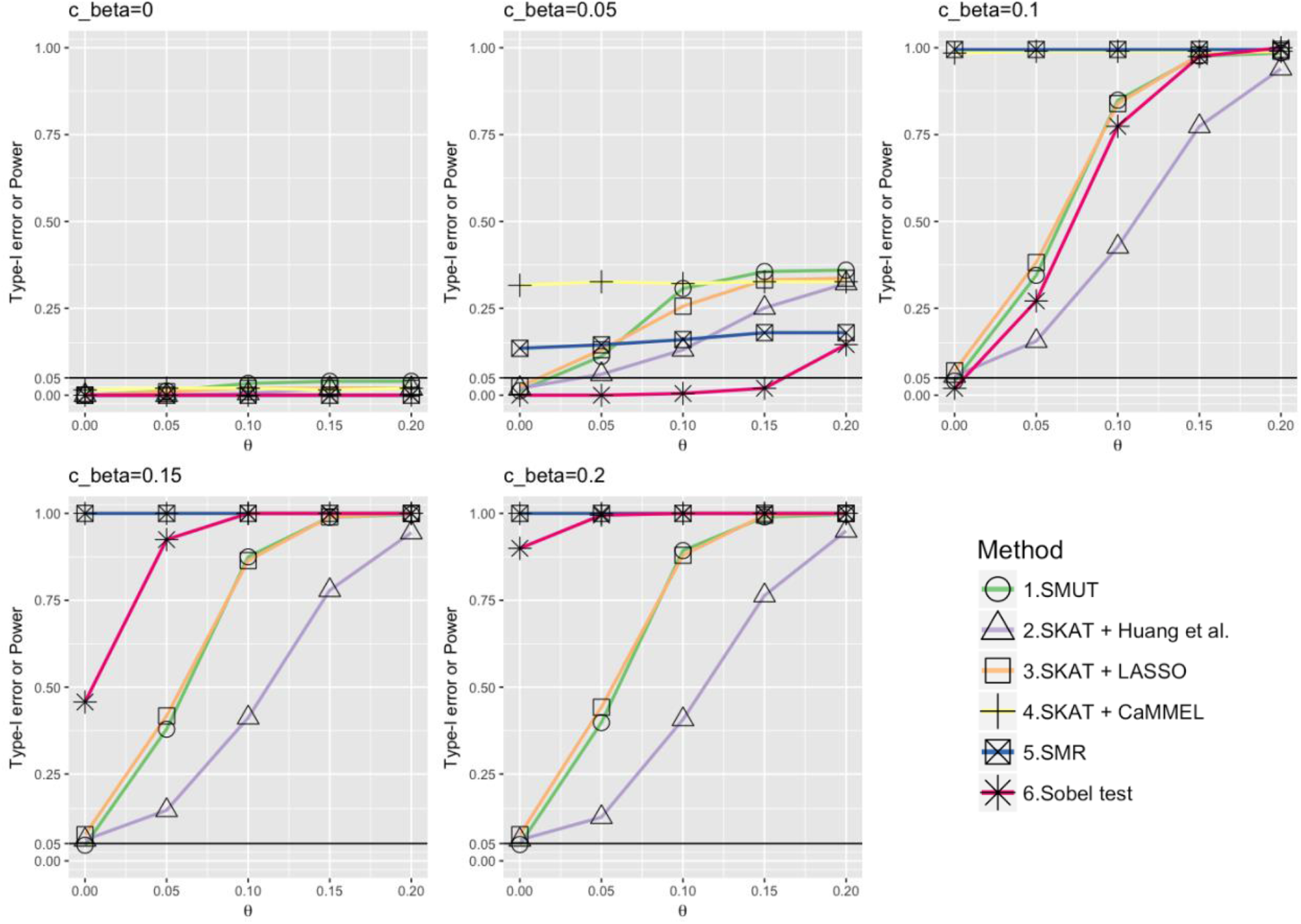
Power and type-I error when causal SNPs are simulated sparse. The x-axis is the true value for mediator’s effect(0) on the outcome in the outcome model. The y-axis is the power or type-I error. The sub-title indicates the true value for *c_β_*. When *c_β_* = 0 or *θ* = 0, it is under the null hypothesis and y-axis represents the corresponding type-I error. When *c_β_* ≠ 0 and *θ* ≠ 0, it is under alternative hypothesis and y-axis represents the corresponding power. The underlying truth for simulated data sets is the sparse scenario in **Table 1**. The candidate SNPs fit in the mediator and outcome model are the 2,891 SNPs with MAF ≥ 1%. The graphs are drawn using the R package ggplot2 (51) and RColorBrewer (52).

**Figure 3.**
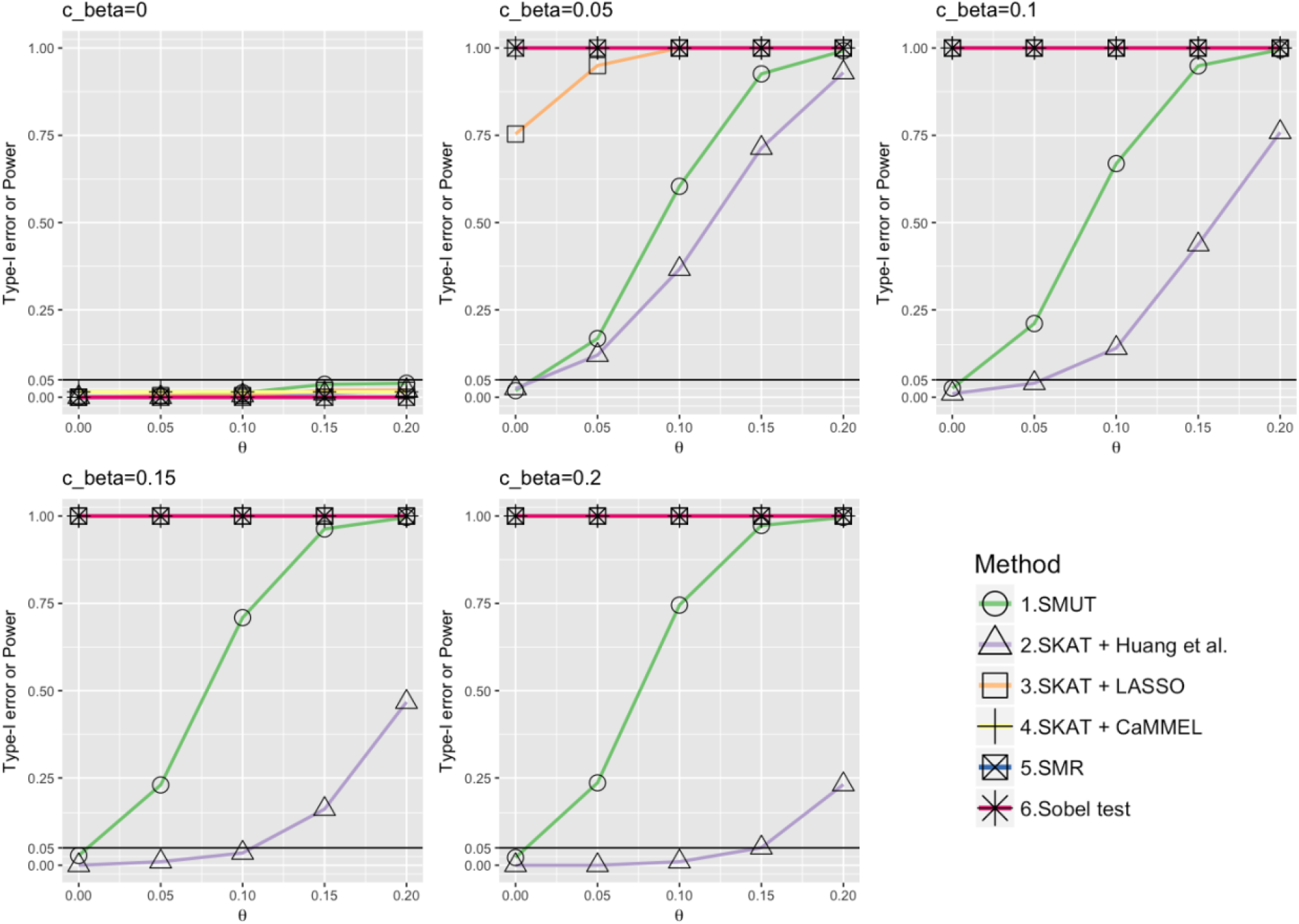
Power and type-I error when causal SNPs are simulated dense. The meaning of x-axis and y-axis is the same as **Figure 2**. The underlying truth for simulated data sets is the dense scenario in **Table 1**. The candidate SNPs fit in the mediator and outcome model are the 2,891 SNPs with MAF ≥ 1%. The graphs are drawn using the R package ggplot2 (51) and RColorBrewer (52).

Assuming normality of *γ_j_*(*j* = 1,2,…,*q*) in the outcome model may not be strictly correct when some SNPs are non-causal (*γ_j_* exactly zero) while others are causal. A mixture distribution would be more appropriate. But our approach gives valid tests in simulations even when the assumption may not be valid.

### Power in Simulations

We assessed power only for tests with protected Type-I error, namely our SMUT and adapted Huang et al. SMUT demonstrated large power gains when the causal SNPs were either sparse or dense. For example, dense causal SNPs when *c_β_* = 0.2, *θ* = 0.15, SMUT and adapted Huang et al. had 97% and 5% power respectively and the power gain was 92%. Power gains appeared more profound with increasing *c_β_*, likely because adapted Huang et al. became very conservative when the pleiotropy effect (*c_β_*) was large.

### Robustness with Alternative Testing Strategies

As aforementioned, the true causal SNPs were drawn from common (MAF > 1%) SNPs and by default all common SNPs were simultaneously modeled and tested. Thus the set of testing SNPs include all the causal SNPs. Alternatively, we considered two other testing strategies: (1) eQTL SNPs only; and (2) SNPs with MAF ≥ 5% only. Under (1), our observations above regarding Type-I error and power remained largely the same: namely SMUT remained valid and more powerful than alternative methods (Figures 4 and 5). In addition, adapted Huang et al. was more powerful using testing strategy (1) than testing all common SNPs in the default setting in most scenarios. For example, with sparse causal SNPs and *c_β_* = 0.05, *θ* = 0.15, adapted Huang et al. had 25% and 96% power using the default and testing strategy (1) while SMUT had 36% and 97% power (Figure 6).

**Figure 4.**
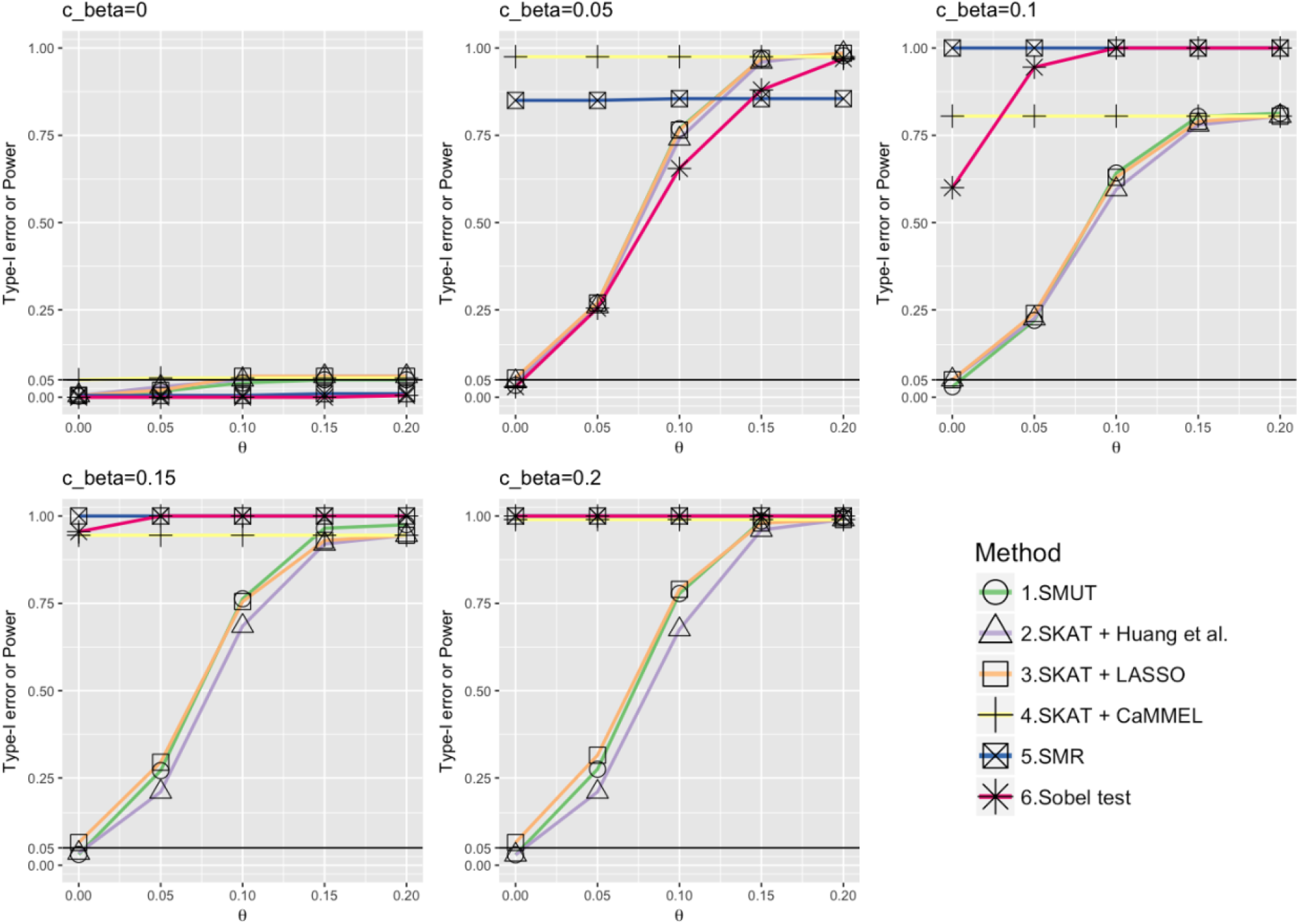
Power and type-I error under alternative setting (1) when testing mediation effects using the true eQTL SNPs and the underlying truth for simulated data sets is the sparse scenario in **Table 1**. The meaning of x-axis and y-axis is the same as **Figure 2**. The graphs are drawn using the R package ggplot2 (51) and RColorBrewer (52).

**Figure 5.**
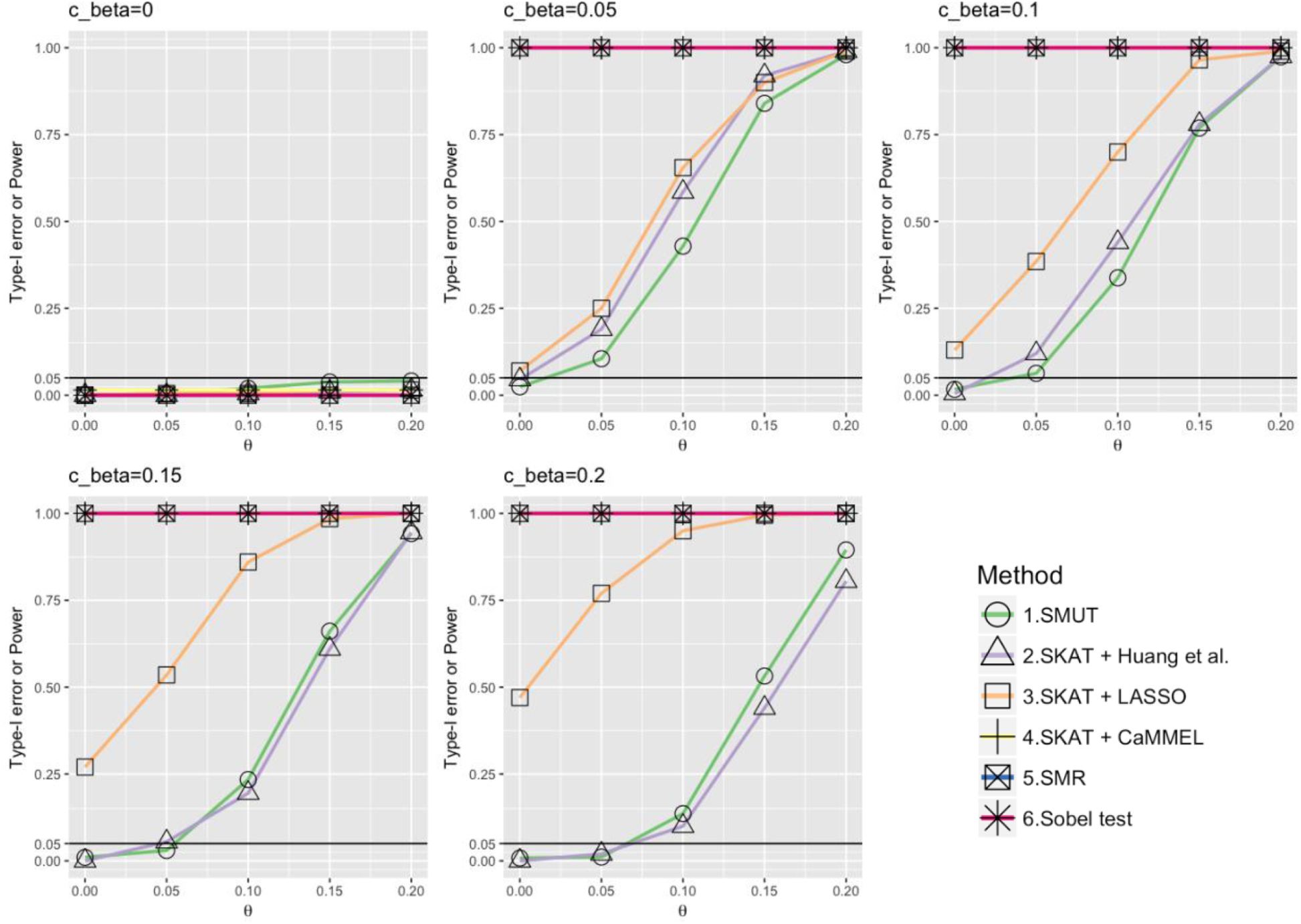
Power and type-I error under alternative setting (1) when testing mediation effects using the true eQTL SNPs and the underlying truth for simulated data sets is the dense scenario in **Table 1**. The meaning of x-axis and y-axis is the same as **Figure 2**. The graphs are drawn using the R package ggplot2 (51) and RColorBrewer (52).

**Figure 6.**
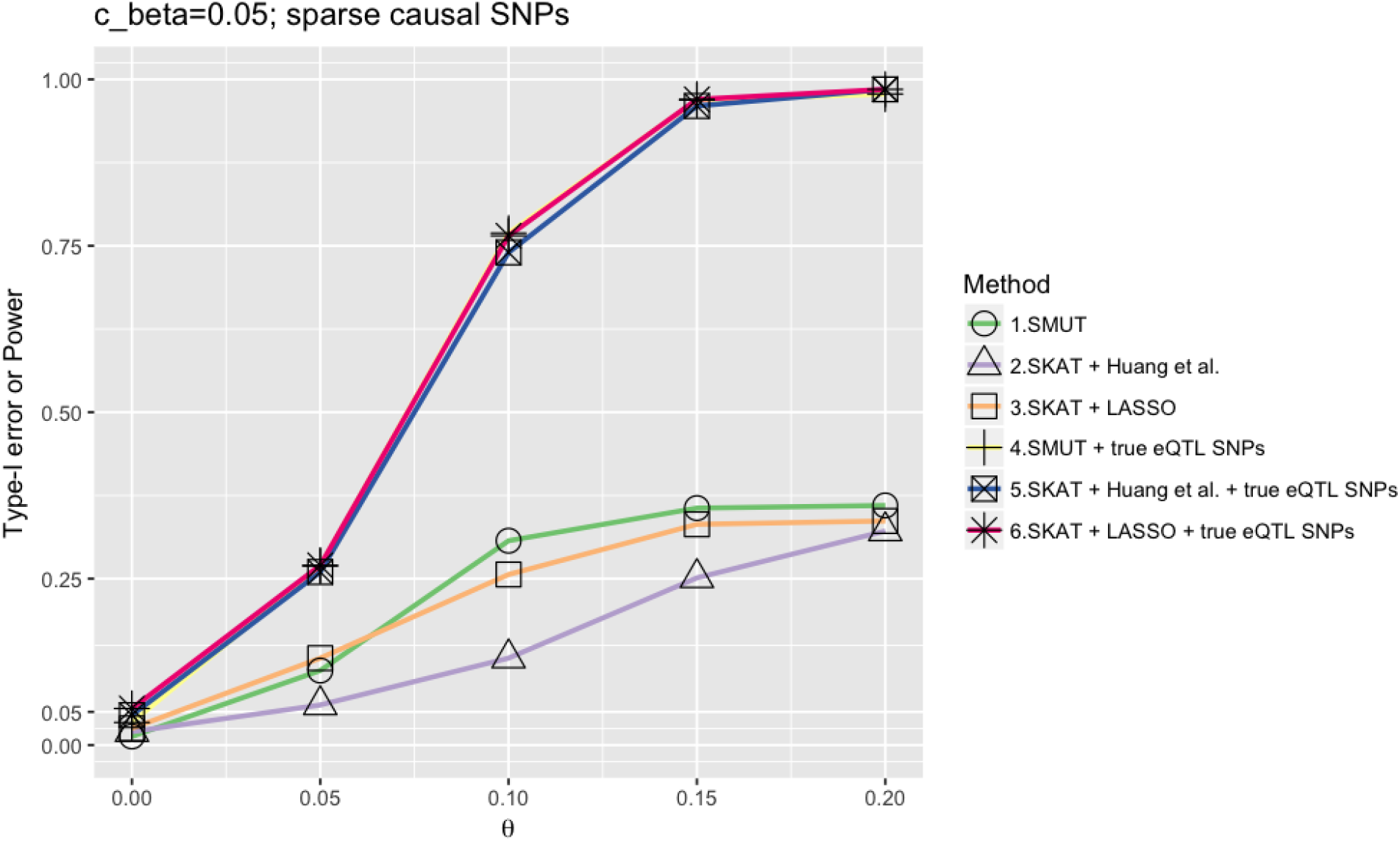
Example of power gain when the true eQTL SNPs are known. Under this situation, with the knowledge on eQTL SNPs helps increase power for SMUT, adaptive LASSO and adaptive Huang et al.’s method. The graphs are drawn using the R package ggplot2 (51) and RColorBrewer (52).

Because SMUT and adapted Huang et al. had protected type-I error, we evaluated only their performance under alternative setting (2). Using testing strategy (2) where only SNPs with MAF ≥ 5% were tested, both SMUT and adapted Huang et al. had inflated type-I error (Figure 7 and 8). This might be due to the violation of confounding assumptions for mediation analysis (31), because shared SNPs became mediator-outcome confounders when absent in models.

**Figure 7.**
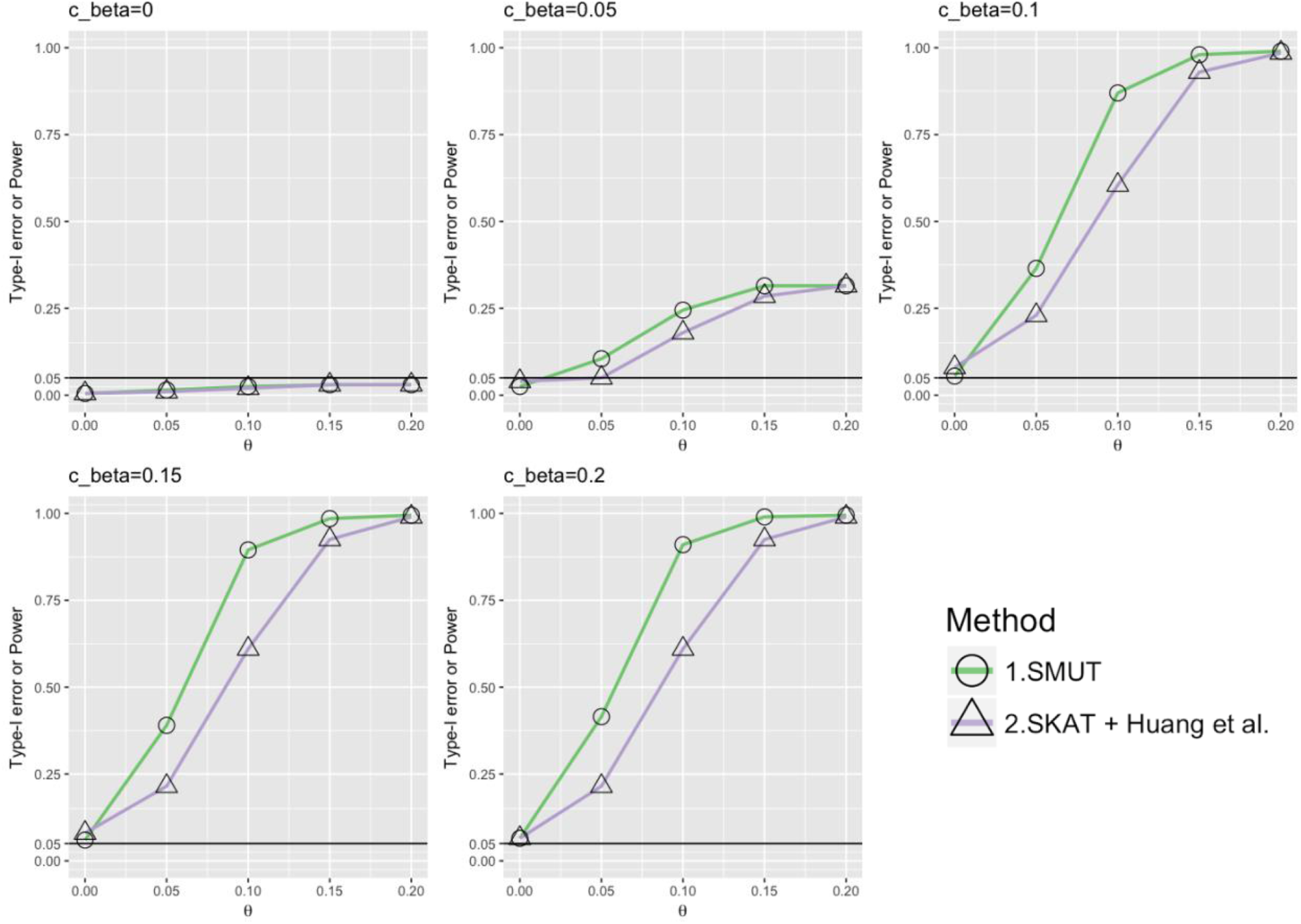
Power and type-I error under alternative setting (2) when testing SNPs with MAF from 1% to 5% and the underlying truth for simulated data sets is the sparse scenario in **Table 1**. The meaning of x-axis and y-axis is the same as **Figure 2**. The graphs are drawn using the R package ggplot2 (51) and RColorBrewer (52).

**Figure 8.**
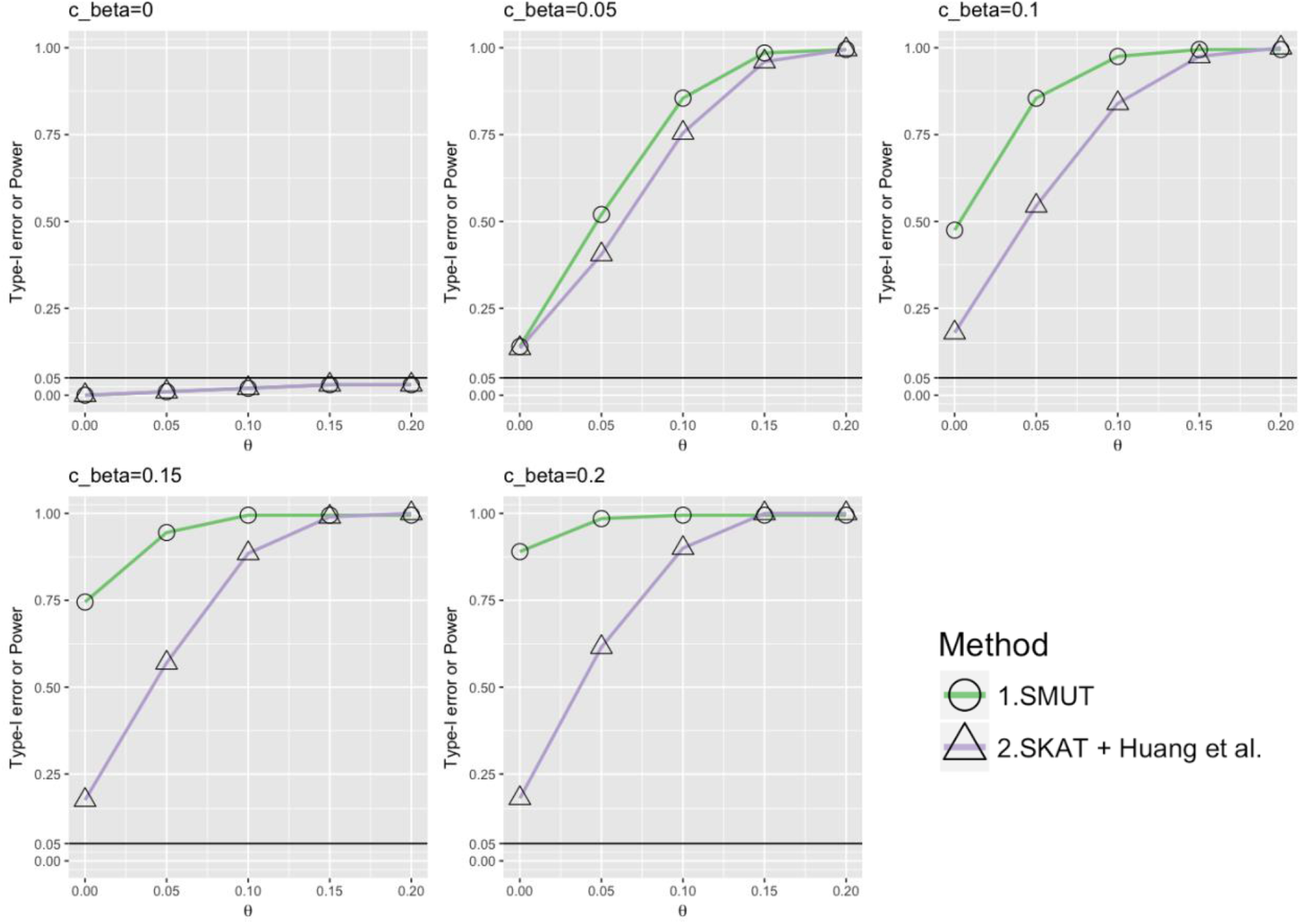
Power and type-I error under alternative setting (2) when testing SNPs with MAF from 1% to 5% and the underlying truth for simulated data sets is the dense scenario in **Table 1**. The meaning of x-axis and y-axis is the same as **Figure 2**. The graphs are drawn using the R package ggplot2 (51) and RColorBrewer (52).

### Real Data Application: METSIM Dataset

The METSIM study is a population-based study with 10,197 males, aged 45–73 years, randomly selected from the population register of Kuopio town in eastern Finland (population 95,000) (45). We analyzed genotype, gene expression and phenotype data in the subset of 770 participants with gene expression measurements from subcutaneous adipose tissue (46). The outcome of interest is plasma adiponectin levels. All METSIM subjects participated in a 1-day outpatient visit to the Clinical Research Unit at the University of Kuopio for data collection, which included an interview for their medical history and a blood sample following a 12-hour fast. Plasma was measured using the Human Adiponectin Elisa kit (LINCO Research).

Here, we tested two “positive control” loci for which our previous study (46) provided mechanistic evidences. The first locus was the adiponectin-associated GWAS locus *ARL15* (with the index SNP rs6450176 being an *ARL15* intronic variant), where the association might be mediated, at least in part, through altered expression of the *FST* gene located further (>521 kb from rs6450176) away instead of *ARL15* (46, 47). The second locus was the *ADIPOQ* locus, also associated with adiponectin levels.

We first extracted SNPs within ±1 Mb of the corresponding genes, *ARL15* union *FST* and *ADIPOQ* union *ADIPOQ-AS1* for the two loci respectively. In terms of phenotypic outcome, namely adiponectin, trait levels were inverse normal transformed after adjusting for age and BMI, following our previous work (46). For the first *ADIPOQ* locus, we tested 286 SNPs with adiponectin association *p* value < 5 × 10^−8^, using SMUT, adaptive Huang et al.’s method, adaptive CaMMEL, CIT, SMR, and Sobel test. Results are summarized in Table 2. Huang et al.’s method returned no results (therefore not shown in Table 2) because it required standardized genotype data which can be undefined for low frequency SNPs. SMUT and SMR both showed significant mediation effects through *ADIPOQ* on adiponectin: SMUT for two probesets and SMR for two probesets. For the second *FST-ARL15* locus, we tested 366 SNPs with MAF ≥ 1% and adiponectin association *p* values < 0.01. Only SMUT detected significant mediation effects through *FST* (but not *ARL15*) on the adiponectin. These results suggest that our SMUT is more powerful for detecting genuine mediation effects.

**Table 2.**
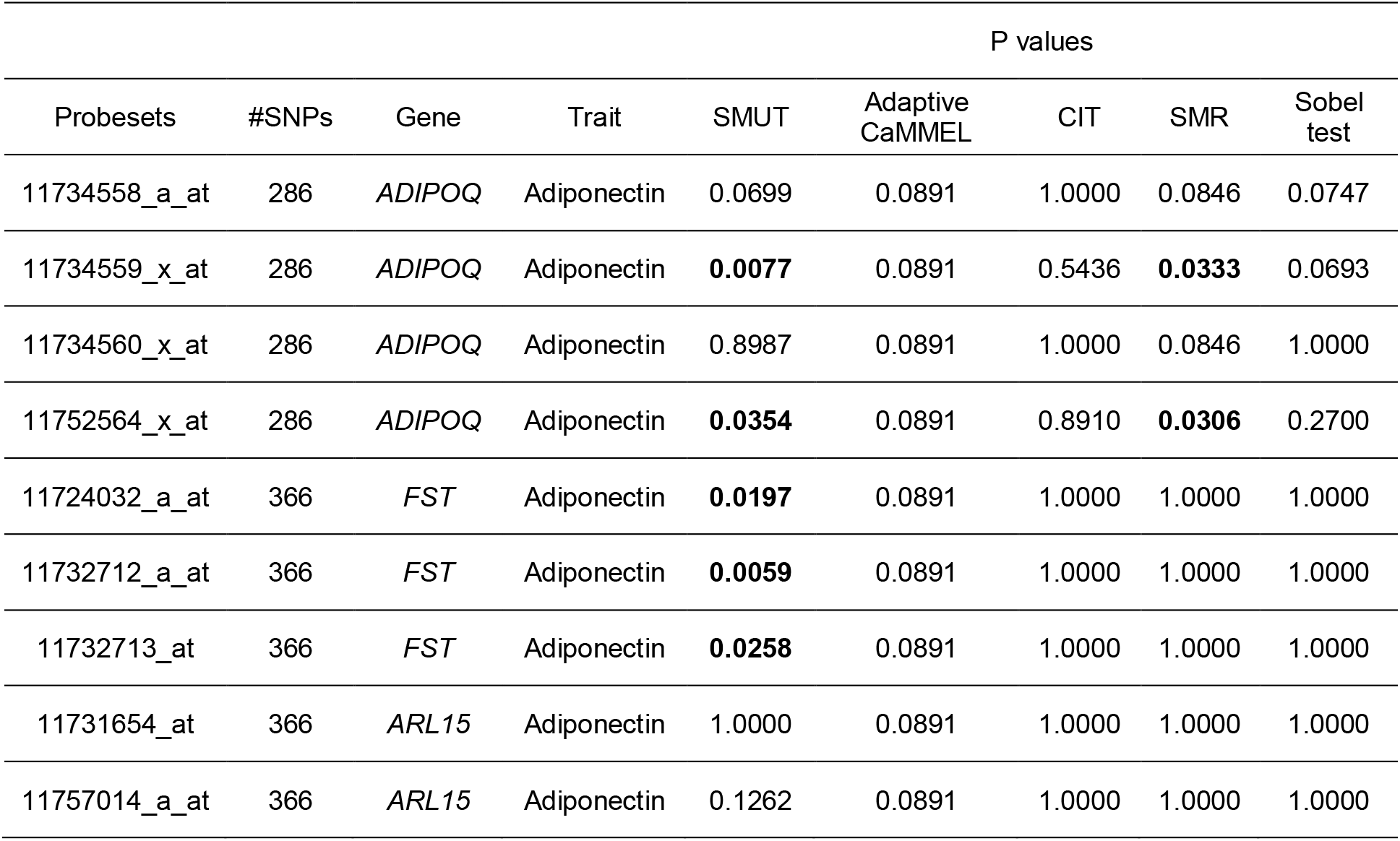
Results from the METSIM study. We used SMUT and other alternative methods (Adaptive CaMMEL, CIT, SMR, Sobel test) to test two loci, the *ARL15* locus and the *ADIPOQ* locus. SNPs within corresponding genes, *ARL15* union *FST* and *ADIPOQ* union *ADIPOQ-AS1* for the two loci respectively, are extracted. For the *ADIPOQ* locus, both SMUT and SMR showed significant mediation effects through *ADIPOQ* on adiponectin. For the *ARL15* locus, only SMUT detected significant mediation effects through *FST* (but not *ARL15*) on the adiponectin. The p values are adjusted using Bonferroni correction.

## DISCUSSION

We propose SMUT, a flexible regression based approach that tests the joint mediation effects of multiple genetic variants on an outcome through a given mediator (e.g. gene). We demonstrate, through extensive simulations, that SMUT preserves type-I error rate and is statistically more powerful than alternative methods including adaptive Huang et al.’s method, adaptive LASSO, adaptive CaMMEL, Sobel test, and SMR.

SMUT has several major advantages over alternative methods. First, as a regression based approach under the mediation analysis framework, SMUT can distinguish mediation from pleiotropy. Second, SMUT generalizes the framework of Baron and Kenny to multiple genetic variants, while methods including SMR and Sobel test can only test one single variant at a time. Third, SMUT naturally accommodates correlation (or LD) among genetic variants while many methods including MR-Egger assume genetic variants under testing are uncorrelated. Finally, SMUT, even its present form, can handle mediators other than gene expression (as presented in the manuscript). For example, molecular measurements such as chromatin spatial organization, histone modification, transcription factor binding affinity, protein abundance can all serve as valid mediators (6, 48–50).

Conceptually, TWAS methods are also designed to elucidate mechanisms regarding the mediation effects of multiple SNPs via gene expression on phenotypic outcome. However, as previously mentioned, TWAS is designed for scenarios where eQTL and GWAS datasets are from two separate sets of study participants. Our SMUT method is designed for the scenario where we have genotype, mediator, and phenotype information measured in the same study subjects. Therefore, we have not directly compared with TWAS methods and deem our SMUT and TWAS useful for different data scenarios.

SMUT can be further extended in several directions. It can be extended to accommodate binary, survival, or longitudinal phenotypic outcome, given its regression based framework. We can also extend SMUT to simultaneously model multiple mediators, which may yield improved power for testing at the price of stronger modelling assumptions.

With more genotyping-based GWAS and large whole genome sequencing efforts underway, the already dauntingly large number of GWAS variants will continue to increase. Approaches generating hypotheses on the mechanisms underlying these variants are imperative. We anticipate SMUT will be a powerful tool in this post-GWAS era to help with bridging the functional gap of GWAS, prioritizing functional follow-up, and disentangling the potential causal mechanism from DNA to phenotype for a new drug discovery and personalized medicine.

## SUPPLEMENTARY DATA

Supplementary Data are available at NAR online.

## ACKNOWLEDGEMENT

We thank the METSIM individuals and investigators to share the data.

## REFERENCES

1. Zhu,X. and Stephens,M. (2017) Bayesian large-scale multiple regression with summary statistics from genome-wide association studies. Ann. Appl. Stat., 11, 1561–1592.

2. Nicolae,D.L., Gamazon,E., Zhang,W., Duan,S., Dolan,M.E. and Cox,N.J. (2010) Trait-associated SNPs are more likely to be eQTLs: annotation to enhance discovery from GWAS. PLoS Genet., 6, e1000888.

3. Emilsson,V., Thorleifsson,G., Zhang,B., Leonardson,A.S., Zink,F., Zhu,J., Carlson,S., Helgason,A., Walters,G.B., Gunnarsdottir,S., et al. (2008) Genetics of gene expression and its effect on disease. Nature, 452, 423.

4. Nica,A.C., Montgomery,S.B., Dimas,A.S., Stranger,B.E., Beazley,C., Barroso,I. and Dermitzakis,E.T. (2010) Candidate causal regulatory effects by integration of expression QTLs with complex trait genetic associations. PLoS Genet., 6.

5. Albert,F.W. and Kruglyak,L. (2015) The role of regulatory variation in complex traits and disease. Nat. Rev. Genet., 16, 197–212.

6. Consortium,Gte. and others (2015) The Genotype-Tissue Expression (GTEx) pilot analysis: Multitissue gene regulation in humans. Science (80-.)., 348, 648–660.

7. Yang,F., Wang,J., Pierce,B.L., Chen,L.S., Aguet,F., Ardlie,K.G., Cummings,B.B., Gelfand,E.T., Getz,G., Hadley,K., et al. (2017) Identifying cis-mediators for trans-eQTLs across many human tissues using genomic mediation analysis. Genome Res.

8. Gamazon,E.R., Wheeler,H.E., Shah,K.P., Mozaffari,S. V., Aquino-Michaels,K., Carroll,R.J., Eyler,A.E., Denny,J.C., Nicolae,D.L., Cox,N.J., et al. (2015) A gene-based association method for mapping traits using reference transcriptome data. Nat. Genet., 47, 1091–1098.

9. Gusev,A., Ko,A., Shi,H., Bhatia,G., Chung,W., Penninx,B.W.J.H., Jansen,R., De Geus,E.J.C., Boomsma,D.I., Wright,F.A., et al. (2016) Integrative approaches for large-scale transcriptome-wide association studies. Nat. Genet., 48, 245.

10. Zhu,Z., Zhang,F., Hu,H., Bakshi,A., Robinson,M.R., Powell,J.E., Montgomery,G.W., Goddard,M.E., Wray,N.R., Visscher,P.M., et al. (2016) Integration of summary data from GWAS and eQTL studies predicts complex trait gene targets. Nat. Genet., 48, 481–487.

11. Mancuso,N., Shi,H., Goddard,P., Kichaev,G., Gusev,A. and Pasaniuc,B. (2017) Integrating gene expression with summary association statistics to identify genes associated with 30 complex traits. Am. J. Hum. Genet., 100, 473–487.

12. Ainsworth,H.F., Shin,S.-Y. and Cordell,H.J. (2017) A comparison of methods for inferring causal relationships between genotype and phenotype using additional biological measurements. Genet. Epidemiol., 41, 577–586.

13. Civelek,M. and Lusis,A.J. (2014) Systems genetics approaches to understand complex traits. Nat. Rev. Genet., 15, 34.

14. Lawlor,D.A., Harbord,R.M., Sterne,J.A.C., Timpson,N. and Smith,G.D. (2008) Mendelian randomization: Using genes as instruments for making causal inferences in epidemiology. Stat. Med., 27, 1133–1163.

15. Davey Smith,G. and Ebrahim,S. (2003) ‘Mendelian randomization’: can genetic epidemiology contribute to understanding environmental determinants of disease? Int. J. Epidemiol., 32, 1–22.

16. Smith,G.D. and Ebrahim,S. (2004) Mendelian randomization: Prospects, potentials, and limitations. Int. J. Epidemiol., 33, 30–42.

17. Katan,M. (1986) Apoupoprotein e isoforms, serum cholesterol, and cancer. Lancet, 327, 507–508.

18. Qi,L. (2009) Mendelian randomization in nutritional epidemiology. Nutr. Rev., 67, 439–450.

19. Burgess,S. and Thompson,S.G. (2015) Mendelian randomization: methods for using genetic variants in causal estimation CRC Press.

20. Bowden,J., Davey Smith,G. and Burgess,S. (2015) Mendelian randomization with invalid instruments: effect estimation and bias detection through Egger regression. Int. J. Epidemiol., 44, 512–525.

21. Barfield,R., Feng,H., Gusev,A., Wu,L., Zheng,W., Pasaniuc,B. and Kraft,P. (2018) Transcriptome-wide association studies accounting for colocalization using Egger regression. Genet. Epidemiol.

22. Solovieff,N., Cotsapas,C., Lee,P.H., Purcell,S.M. and Smoller,J.W. (2013) Pleiotropy in complex traits: Challenges and strategies. Nat. Rev. Genet., 14, 483–495.

23. Zhu,X., Stephens,M. and others (2017) Bayesian large-scale multiple regression with summary statistics from genome-wide association studies. Ann. Appl. Stat., 11, 1561–1592.

24. Bowden,J., Smith,G.D. and Burgess,S. (2015) Mendelian randomization with invalid instruments: Effect estimation and bias detection through Egger regression. Int. J. Epidemiol., 44, 512–525.

25. Millstein,J., Zhang,B., Zhu,J. and Schadt,E.E. (2009) Disentangling molecular relationships with a causal inference test. BMC Genet., 10.

26. Huang,Y.-T., VanderWeele,T.J. and Lin,X. (2014) Joint analysis of SNP and gene expression data in genetic association studies of complex diseases. Ann. Appl. Stat., 8, 352–376.

27. Huang,Y.T. (2015) Integrative modeling of multi-platform genomic data under the framework of mediation analysis. Stat. Med., 34, 162–178.

28. Huang,Y.-T., Liang,L., Moffatt,M.F., Cookson,W.O.C.M. and Lin,X. (2015) iGWAS: Integrative Genome-Wide Association Studies of Genetic and Genomic Data for Disease Susceptibility Using Mediation Analysis. Genet. Epidemiol., 39, 347–356.

29. Lin,X. (1997) Variance component testing in generalised linear models with random effects. Biometrika, 84, 309–326.

30. MacKinnon,D.P., Fairchild,A.J. and Fritz,M.S. (2007) Mediation Analysis. Annu. Rev. Psychol., 58, 593–614.

31. VanderWeele,T.J. (2016) Mediation Analysis: A Practitioner’s Guide. Annu. Rev. Public Health, 37, 17–32.

32. Sobel,M.E. (1982) Asymptotic confidence intervals for indirect effects in structural equation models. Sociol. Methodol., 13, 290–312.

33. Sobel,M.E. (1986) Some new results on indirect effects and their standard errors in covariance structure models. Sociol. Methodol., 16, 159–186.

34. Baron,R.M. and Kenny,D. a (1986) The moderator-mediator variable distinction in social psychological research: conceptual, strategic, and statistical considerations. J. Pers. Soc. Psychol., 51, 1173–1182.

35. Berger,R.L. and Hsu,J.C. (1996) Bioequivalence trials, intersection-union tests and equivalence confidence sets. Stat. Sci., 11, 283–319.

36. Wu,M.C., Lee,S., Cai,T., Li,Y., Boehnke,M. and Lin,X. (2011) Rare-variant association testing for sequencing data with the sequence kernel association test. Am. J. Hum. Genet., 89, 82–93.

37. Lee,S., Abecasis,G.R., Boehnke,M. and Lin,X. (2014) Rare-variant association analysis: study designs and statistical tests. Am. J. Hum. Genet., 95, 5–23.

38. Radhakrishna Rao,C. and Bartlett,M.S. (1948) Large sample tests of statistical hypotheses concerning several parameters with applications to problems of estimation. Math. Proc. Cambridge Philos. Soc., 44, 50.

39. Engle,R.F. (1984) Wald, likelihood ratio, and Lagrange multiplier tests in econometrics. Handb. Econom., 2, 775–826.

40. Dempster,A.P., Laird,N.M. and Rubin,D.B. (1977) Maximum Likelihood from Incomplete Data via the EM Algorithm. J. R. Stat. Soc. Ser. B, 39, 1–38.

41. McCulloch,C.E., Searle,S.R. and Neuhaus,J.M. (2008) Generalized, Linear, and Mixed Models, 2nd Edition.

42. Schaffner,S.F., Foo,C., Gabriel,S., Reich,D., Daly,M.J. and Altshuler,D. (2005) Calibrating a coalescent simulation of human genome sequence variation. Genome Res., 15, 1576–1583.

43. Tibshirani,R. (1996) Regression shrinkage and selection via the lasso. J. R. Stat.Soc. Ser. B.

44. Zhao,P. and Yu,B. (2006) On Model Selection Consistency of Lasso. J. Mach.Learn. Res., 7, 2541–2563.

45. Stancakova,A., Javorsky,M., Kuulasmaa,T., Haffner,S.M., Kuusisto,J. and Laakso,M. (2009) Changes in Insulin Sensitivity and Insulin Release in Relation to Glycemia and Glucose Tolerance in 6,414 Finnish Men. Diabetes, 58, 1212–1221.

46. Civelek,M., Wu,Y., Pan,C., Raulerson,C.K., Ko,A., He,A., Tilford,C., Saleem,N.K., Stančáková,A., Scott,L.J., et al. (2017) Genetic Regulation of Adipose Gene Expression and Cardio-Metabolic Traits. Am. J. Hum. Genet., 100, 428–443.

47. Martin,J.S., Xu,Z., Reiner,A.P., Mohlke,K.L., Sullivan,P., Ren,B., Hu,M. and Li,Y. (2017) HUGIn: Hi-C unifying genomic interrogator. Bioinformatics, 33, 3793–3795.

48. Schmitt,A.D., Hu,M., Jung,I., Xu,Z., Qiu,Y., Tan,C.L., Li,Y., Lin,S., Lin,Y., Barr,C.L., et al. (2016) A Compendium of Chromatin Contact Maps Reveals Spatially Active Regions in the Human Genome. Cell Rep., 17, 2042–2059.

49. Xu,Z., Zhang,G., Jin,F., Chen,M., Furey,T.S., Sullivan,P.F., Qin,Z., Hu,M. and Li,Y. (2015) A hidden Markov random field-based Bayesian method for the detection of long-range chromosomal interactions in Hi-C data. Bioinformatics, 32, 650–656.

50. Sun,W., Kechris,K., Jacobson,S., Drummond,M.B., Hawkins,G.A., Yang,J., Chen,T., Quibrera,P.M., Anderson,W., Barr,R.G., et al. (2016) Common Genetic Polymorphisms Influence Blood Biomarker Measurements in COPD. PLoS Genet, 12, 1–33.

51. Wickham,H. (2016) ggplot2: elegant graphics for data analysis Springer.

52. Harrower,M. and Brewer,C.A. (2003) ColorBrewer. org: an online tool for selecting colour schemes for maps. Cartogr. J., 40, 27–37.

